# Effects of Ocean Climate on Spatiotemporal Variation in Sea Urchin Settlement and Recruitment

**DOI:** 10.1101/387282

**Authors:** Daniel K. Okamoto, Stephen Schroeter, Daniel C. Reed

## Abstract

Sea urchins are voracious herbivores that influence the ecological structure and function of nearshore ecosystems throughout the world. Like many species that produce planktonic larvae, their recruitment is thought to be particularly sensitive to climatic fluctuations in temperature that directly or indirectly affect adult reproduction and larval transport and survival. Yet how climate alters sea urchin populations in space and time by modifying larval recruitment and year-class strength on the time-scales that regulate populations remains understudied. Using a, spatially replicated weekly-biweekly dataset spanning 27 years and 1100 km of coastline, we characterized seasonal, interannual, and spatial patterns of larval settlement of the purple sea urchin (*Strongylocentrotus purpuratus*). We show that large spatial differences in temporal patterns of larval settlement were associated with different responses to fluctuations in ocean temperature and climate. Importantly, we found a strong correlation between larval settlement and regional year class strength suggesting that such temporal and spatial variation in settlement plays an important role in controlling population dynamics. These results provide strong evidence over extensive temporal and spatial domains that climatic fluctuations shape broad-scale patterns of larval settlement and subsequent population structure of an important marine herbivore known to control the productivity, community state and provisioning services of marine ecosystems.

## Introduction

Large scale climate oscillations (e.g., El Niño/Southern Oscillation, North Atlantic Oscillation) lead to changes in ocean temperature, biogeochemistry and the severity and frequency of disruptive events that affect ocean circulation, upwelling and primary productivity (2014; Mantua et al., 1997). Such shifts impose wide-reaching ecological impacts, in part by altering animal recruitment and food web structure in space and time (Sydeman et al., 2015). Hence, understanding how climate variability alters the recruitment of marine species is particularly important for effective conservation and management of the ocean’s resources. Climatic fluctuations give rise to shifts in numerous factors that shape both adult reproduction and larval supply, including primary productivity, temperature, and advection and transport. Given these multiple direct and indirect effects of climate on recruitment, significant challenges remain in achieving such understanding for benthic species with planktonic larvae due to the substantial effort needed to characterize spatial and temporal variation in larval settlement and the numerous sensitive vital rates that contribute to it.

For benthic species like sea urchins, understanding causes and consequences of recruitment variability has both ecological and economic implications. Sea urchin grazing can alter the structure of some of the world’s most diverse and productive marine ecosystems, including coral reefs (Edmunds & Carpenter, 2001), seagrass meadows (reviewed by Valentine & Heck Jr, 1999) and kelp forests (reviewed by Filbee-Dexter & Scheibling, 2014). In addition, sea urchins form the basis of important nearshore fisheries in many regions of the world (e.g. Andrew et al., 2003; Kato & Schroeter, 1985). As a result, climate-driven changes in sea urchin populations have the potential to profoundly affect the ecological structure and functioning of marine ecosystems and the economic value of the fisheries that they support. Much of the research on controls of sea urchin population dynamics has focused on the roles of predation and disease in controlling adult abundance and their cascading influence on community structure (e.g. Burt et al., 2018; Estes & Duggins, 1995; Filbee-Dexter & Scheibling, 2014; Lafferty, 2004). Yet short-term empirical studies (months to a few years) suggest that environmentally regulated larval supply is likely an important driver of adult urchin dynamics (Hernández et al., 2010; Ling et al., 2009). Despite the widespread recognition of the importance of recruitment variation in controlling population fluctuations in many marine species (Shelton & Mangel, 2011), relatively few studies have examined the biotic and abiotic processes controlling the supply of sea urchin larvae in nature (but see Hernández et al., 2010; Ling et al., 2009), and the degree to which they affect the abundance and dynamics of older life stages over time scales that impact population dynamics.

Fluctuations in climate can affect spatial and temporal patterns of larval supply by influencing the production of larvae by benthic adults, transport of larvae to adjacent habitats, and the survival of larvae in the plankton. Increases in ocean temperature can impact larval production by: (1) increasing adult mortality via the spread of water-borne pathogens (reviewed by Feehan & Scheibling, 2014), and (2) reducing adult fecundity and inhibiting gametogenesis by altering food quantity and quality (Basch & Tegner, 2007; Cochran & Engelmann, 1975; Foster et al., 2015; Okamoto, 2014). Because sea urchins produce feeding larvae that spend weeks to months in the plankton, increases in ocean temperature can also affect larval development, growth and survival, either directly, or indirectly by altering the availability of their phytoplankton food source (Bertram & Strathmann, 1998; Byrne et al., 2009; Hoegh-Guldberg & Pearse, 1995; Strathmann, 1987). Finally, climate related changes in patterns of ocean circulation can affect the transport of larvae from source to destination (but see Morgan, 2014; Siegel et al., 2008). Thus, the effects of climatic change on sea urchin recruitment represent cumulative impacts on adult abundance and reproduction, complex current patterns that transport larvae, behavioral responses of larvae, and larval development and survival. Because patterns of ocean temperature, circulation and upwelling can vary dramatically in space, the effects of climate oscillations on sea urchin recruitment potentially vary over large spatial scales. A dearth of long-term, high frequency, spatially extensive data has prevented characterizing temporal and spatial variability in larval settlement in sea urchins, the degree to which it is explained by different sources of environmental variation, and the relative importance of these drivers in accounting for fluctuations in population size.

Here we analyzed a 27-year weekly to biweekly time series of the recruitment of newly metamorphosed larvae (hereafter referred to as larval settlement) at sites distributed across 1100 km of coast in California to investigate sources of spatial and temporal variability in larval settlement of the purple sea urchin *Strongylocentrotus purpuratus*. Our objectives were to: (1) quantify variation in larval settlement across different temporal and spatial scales; (2) evaluate whether larval settlement on artificial substrates predicts year-class strength in natural populations; and (3) determine the relative importance of adult abundance, larval and adult food supply, sea surface temperature and broad-scale fluctuations in ocean climate in contributing to the observed variation in larval settlement

### Study system

Populations of the purple sea urchin (*Strongylocentrotus purpuratus*) occupy shallow subtidal and intertidal rocky substrata from at least 27°N on the western coast of the Baja Peninsula (Olivares-Bañuelos et al., 2008) to at least 59°N on the Kenai Peninsula in Alaska (Field & Walker, 2003). Purple urchins are broadcast spawners and the seasonality of their spawning is generally thought to be driven by spring photoperiod and temperature (Cochran & Engelmann, 1975; Gonor, 1973; Pearse et al., 1986). Energy available for gonad production is largely determined by macroalgal food availability; specifically, sea urchins in barrens devoid of abundant macroalgae can show dramatic reductions in fecundity (>99%), and resumption of feeding in emaciated urchins can lead to gonadal recovery within 2-3 months (Okamoto, 2014). Results from the field and laboratory indicate that the thermal upper limit to completion of gametogenesis is about 17°C (Basch & Tegner, 2007; Cochran & Engelmann, 1975). Fertilized zygotes develop into planktonic echinoplutei larvae that feed exclusively on phytoplankton (Strathmann, 1987). After spending several weeks to months in the plankton, individuals begin final metamorphosis and settle to the benthos (Strathmann, 1978) at a size of ∼500 μm in diameter (Okamoto unpublished data). Larval settlement varies dramatically among locations at both small and large spatial scales (Ebert, 2010). Once settled, fully competent individuals occupy cobble and other complex substrata and between 12-24 months become visible in benthic surveys at 1-2 cm diameter. For a full review see Rogers-Bennet & Okamoto (Rogers-Bennett & Okamoto, 2020).

### Collection of newly settled urchins along the coast of California, USA (1990-2016)

Settlement of newly metamorphosed purple sea urchins was sampled in three major regions along the California coast from 1990 through 2016, with a total of 54,588 replicate observations. Sampling regions (from south to north) included two sites in San Diego County (Scripps Pier and Ocean Beach Pier), four sites in the Santa Barbara Channel (Anacapa Island, Stearns Wharf, Ellwood Pier and Gaviota Pier) and one site at Point Cabrillo in Fort Bragg (Figure 1). San Diego and the Santa Barbara Channel lie within the Southern California Bight and Fort Bragg is in northern California. At each site, newly settled urchins were collected using nylon-bristled scrub brushes (2.5 cm long bristles and a 6 x 9 cm wooden base) suspended 1 to 2 m from the benthos (Ebert et al., 1994). The majority of deployments included 4-8 replicate brushes collected weekly at each site from 1990 to 2003, and biweekly thereafter through 2016. Upon collection brushes were transported to the laboratory in plastic bags and rinsed through a 350 μm mesh sieve. Purple urchins were sorted from other species, counted and preserved. A more detailed description of the sampling methods, geographic and taxonomic coverage and links to the data and metadata are provided in Schroeter et al. (2019)

**Figure 1:**
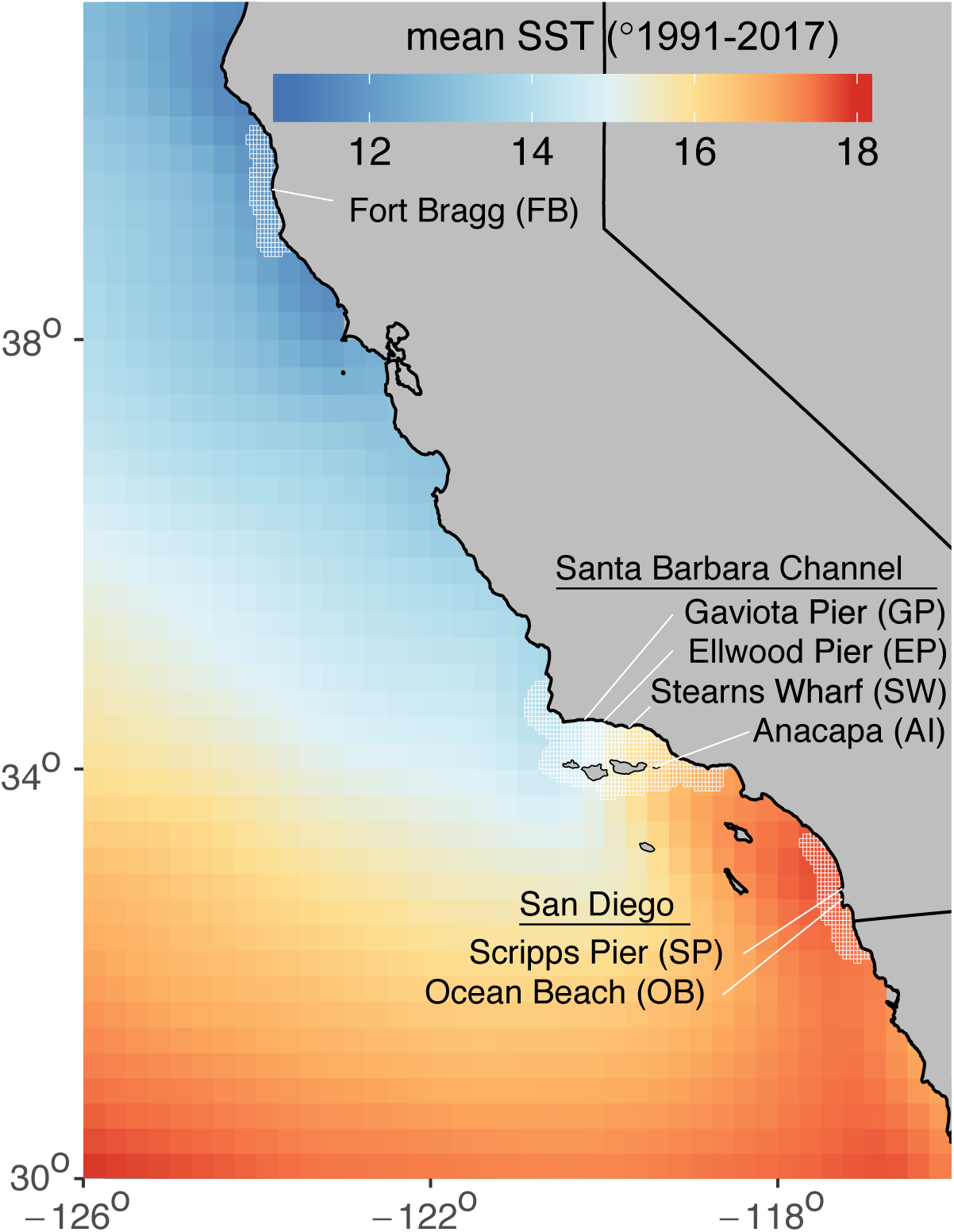
Map of larval settlement collection sites in California. Colors represent the average spatial gradient of sea surface temperature from 1991-2017. The hatched white lines show the shoreline buffer used to constrain sea surface temperature and sea surface chlorophyll index to local larval settlement observations.

## Methods

### Spatio-temporal trends in larval settlement

We estimated seasonal, annual, and spatial patterns in larval settlement using an integrated spatiotemporal Bayesian model that accounted for the spatially and temporally intercorrelated and heterogeneous nature of the multivariate time series. We used a Bayesian state-space formulation for the model for several reasons. First, episodic periods of low replication, low observation numbers, missing data, or slight variation in the sampling interval mean that the true number of settlers may not be reflected in the empirical mean value from brush data (i.e. the true value is not always observable). Second, when brushes contained hundreds or more individual urchins, species identification consisted of subsampling urchins to estimate proportions of individual species (in this case *S. purpuratus* versus the rarer red urchin, *Mesocentrotus franciscanus*). This subsampling routine requires accounting for the uncertainty in the sampled species ratios in the estimation of settler abundance, which we did by incorporating a Bayesian prior in the form of a Beta distribution with hyperparameters as the observed ratio and total subsample count (the Jeffrey’s prior for ratios from binomial counts). Third, initial examination of the data indicated substantial seasonal and interannual variability that was multiplicative among-samples within each period, as well as serial autocorrelation and spatial components that needed to be simultaneously accounted for. We used a Poisson observation likelihood to link the biweekly mean estimate from the trend equation (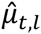 - see Table 1, Equations 3 & 4, and Table 2) to observed counts of larval settlers (N) for each brush (b) within each site within each year. To account for the above described nuances of the data we estimated biweekly settlement density 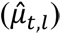 as the sum of a site-specific mean (β_0,*l*_), log annual trend (Annual_Trend*_s,t_*), log seasonal trend (Seas_Trend*_t,l_*), and a spatially correlated lognormal process error (ε*_t,l_*);

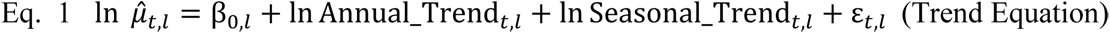

**Table 1:**
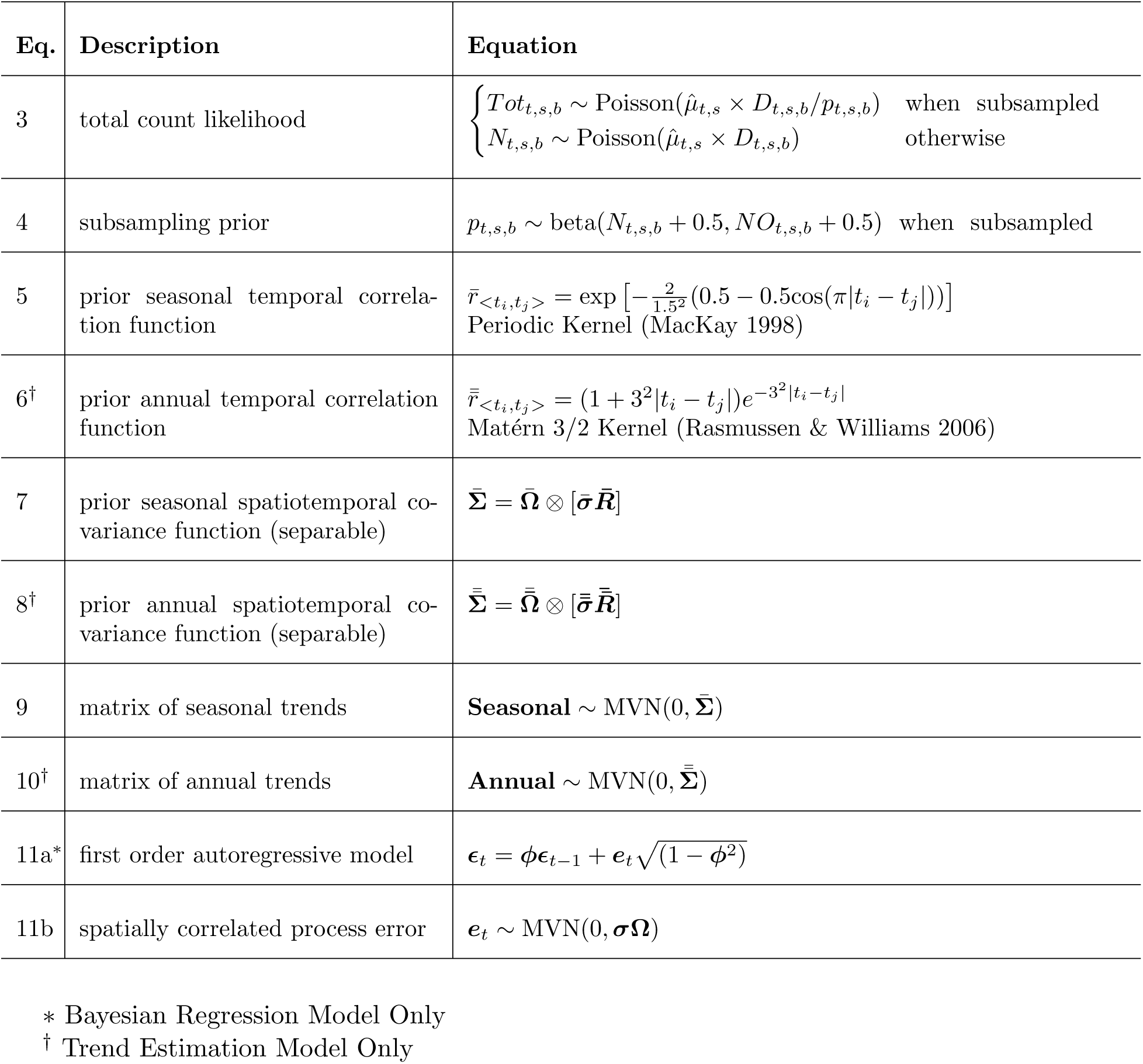
Equations for the multivariate spatiotemporal models. Variables, parameters and priors are described in Table 2. Note that there are two primary model forms: a trend estimation model, and a Bayesian regression model. Equations apply to both model forms except where specified.

**Table 2:** Definition of parameters, variables, and priors for equations described in Table 1

| Symbol | Description | Type | Prior/Input |
| --- | --- | --- | --- |
| $\hat{\mu}_{t,s}$ | expected settlement density (per brush, per day) of purple urchins, defined in Eqs. 1 or 2 in the main text for each model. | estimated | Eqs. 1, 2 |
| $N_{t,s,b}$ | number of purple urchins counted at time $t$, site $s$ and brush $b$. When urchin species within brushes are subsampled, data produced are a total urchin count ($Tot_{t,s,b}$), a total subsample count ($NO_{t,s,b}$), and a purple urchin subsample count ($N_{t,s,b}$). | input | data |
| $Tot_{t,s,b}$ | total number of urchins counted at time $t$, site $s$ and brush $b$ | input | data |
| $p_{t,s,b}$ | estimated proportion of purple urchins in the counts given the subsample (when urchin counts in brushes are subsampled). | input | data |
| $D_{t,s,b}$ | exact number of days of the brush deployment | input | data |
| $\mathbf{X}$ | covariate design matrix - for estimation model represents only site indices, for regression model includes covariates | input | data |
| $\beta$ | vector of site specific scale parameters and covariate coefficients | estimated | **Scale parameters:**<br>Student-t: scale=2;<br>**Coefficients:**<br>regularized horseshoe prior (see methods). |
| $\phi$ | vector of site specific first order autoregressive parameters & estimated (regression model only) | estimated | Uniform -0.95 to 0.95 |
| $\Omega, \bar{\Omega}, \bar{\bar{\Omega}}$ | spatial correlation matrices for process error, annual trends, and seasonal trends (estimated through the cholesky factor) | estimated | LKJ prior:scale= 2<br>(Lewandowski *et al.* 2009) |
| $\bar{R}, \bar{\bar{R}}$ | temporal correlation matrices for annual and seasonal gaussian processes | input | eqs 5 & 6 |
| $\sigma, \bar{\sigma}, \bar{\bar{\sigma}}$ | site specific standard deviation for process error, annual trends, and seasonal trends | estimated | half-Cauchy: scale = 2.5<br>(Polson & Scott 2012) |

We estimated trends on the log-scale so that process errors, and seasonal and interannual trends were multiplicative and strictly positive on the original scale.

The annual trends were assumed to be correlated in both space and time, where spatial and temporal covariances were assumed to be independent. The annual spatial covariance was assumed to be unstructured (i.e. no formal distance decay structure) because we only had seven sites. The annual temporal covariance was determined by a Matérn 3/2 kernel because it provided sufficient flexibility to capture interannual trends. The seasonal trend (log-scale) within each site was estimated using a seasonal periodic temporal kernel within the model to capture the cyclical nature of seasonality. The process error (ε*_t,l_*) was modeled with an unstructured spatial covariance to account for correlations among sites in their deviations from model expectations. We estimated the model posteriors using Stan (Carpenter et al., 2016) with three 1000 iteration chains after a 1000 iteration burn-in. Stan model code is provided in the supplement. For model equations and symbology see Tables 1 and 2.

### Relationship between benthic year-class strength and larval settlement

If year class strength of sea urchins is limited by larval supply, then we should see increases (decreases) in the abundance of juveniles in years following anomalously high (low) larval settlement. We therefore tested whether larval settlement in the Santa Barbara Channel (mean of Anacapa, Stearns Wharf, Ellwood Pier and Gaviota Pier) was predictive of subsequent juvenile recruitment (hereafter referred to as “year-class strength”) on natural reefs in the region. We calculated densities of juveniles (<2cm test diameter) from the Channel Islands Kelp Forest Monitoring (KFM) Program (Kushner et al., 2013), including only the 9 sites with time series extending from 1990 through 2016. We examined the relationship between larval settlement and year-class strength using a generalized linear mixed effects model (GLMM) with a negative binomial likelihood (a Poisson likelihood indicated overdispersion) and survey site as a random effect. We tested the hypothesis of no correlation using a likelihood ratio test comparing the models that do versus do not include settlement as a covariate. Models were estimated using glmmTMB (Magnusson et al., 2017) in R (see Electronic Supplement for further details).

### Relationships between larval settlement and biotic and abiotic conditions

We estimated the strength of relationships between biweekly larval settlement and various hypothesized physical and biological drivers using an integrated regression model that included a regularized regression within a Bayesian time-series modelling framework (Fig. 2). We used a Poisson observation likelihood to link estimated latent trends 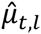 to the settlement count observations (see Table 1 Equations 3 & 4, and Table 2). We estimated the log biweekly density 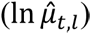 as a function of the centered and standardized covariates (denoted by the vector ***x****_t,l_*) while directly estimating and accounting for seasonal trends (Seasonal_Trend*_t,l_*), and a separable spatially and temporally correlated, multivariate lognormal process error (ε*_t,l_*):

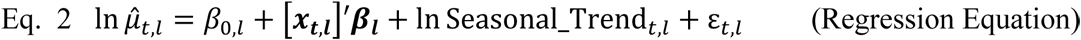

**Figure 2:**
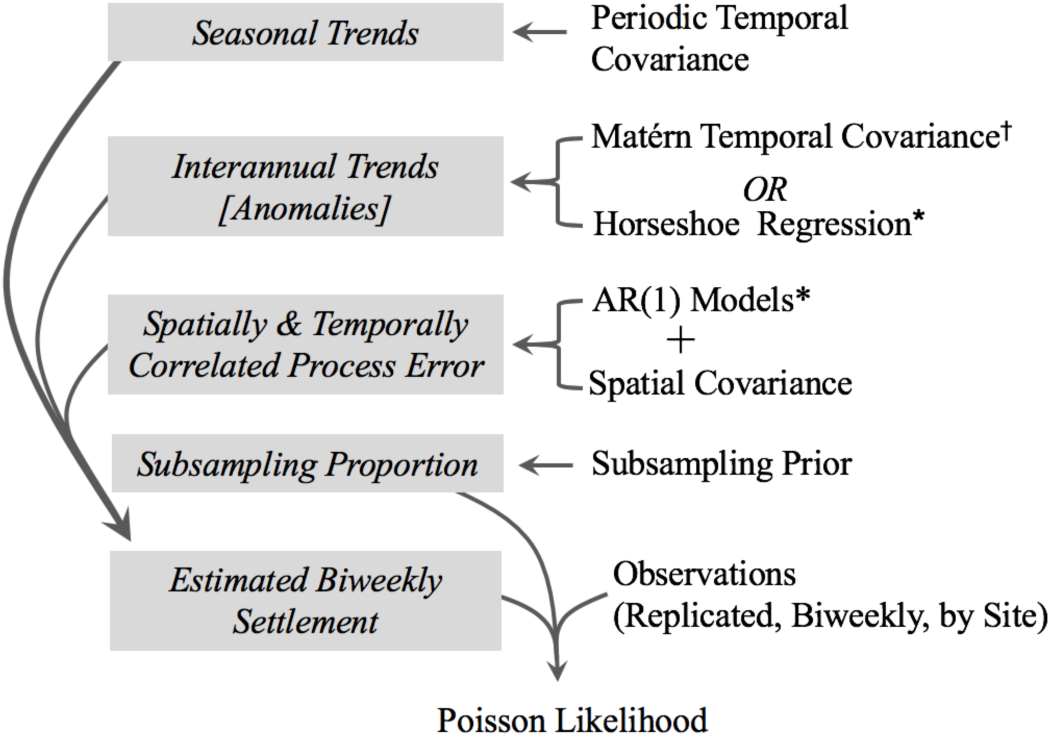
Diagram of the model structure using either a Matérn temporal covariance (trend estimation model) or a horseshoe regression (correlation estimation model) to estimate seasonal anomalies. For details see Tables 1 and 2. *Bayesian Regression Model Only. ^†^Trend Estimation Model Only

We assumed the temporal correlation in the process error ε*_t,l_* followed a first order autoregressive model.

Because these analyses were correlative, we focused on unbiased parameter estimation rather than hypothesis testing or model comparison per se. We estimated two separate models: one included all locally measured (i.e. for a given site) environmental variables and the other included a common set of composite indices of oceanographic climate (“global covariates” – ENSO Index, Pacific Decadal Oscillation, North Pacific Gyre). Because inclusion of numerous covariates can cause overfitting that leads to bias and uncertainty in the explanatory power of the covariates, we erred on the side of sparsity and assigned the vector of regression coefficients (**β*_l_***) a regularized horseshoe prior (i.e. the Finnish Horseshoe). Sparsity in this case assumed that only a few of the covariates were meaningful without a priori knowledge of which covariates were relevant and which were not. A sparse regression encoded this assumption to allow the data to inform which covariates were relevant and how they were correlated with the response variable. The Finnish horseshoe prior is a Bayesian version of the lasso (Carvalho et al., 2010; Piironen & Vehtari, 2017) that produced a data-driven reduction in the influence of weaker covariates by regularization of those coefficients given the data. For full details and model equations, see Tables S.1 and S.2.

Below are the covariates used in the integrated regression model:

#### (i) Oceanographic climate indices (monthly, 1990-2016, all sites)

We used three major global indices of oceanographic climate. The multivariate El Niño Southern Oscillation Index (MEI), the Pacific Decadal Oscillation (PDO - Mantua & Hare, 2002) and the North Pacific Gyre Oscillation (NPGO - Di Lorenzo et al., 2008). The MEI provides a metric of the intensity of El Niño Southern Oscillation (ENSO) fluctuations, which persist for 6-18 months and explains much of the oceanographic variability in the tropics (Di Lorenzo et al., 2013; Wolter & Timlin, 1993; Wolter & Timlin, 1998). In contrast, the PDO and NPGO exhibit longer-period fluctuations, explain much of the oceanographic variability in higher latitudes, and the low frequency variability of these metrics are shaped through ENSO forcing of the Aleutian Low and the North Pacific Oscillation, respectively (Di Lorenzo et al., 2013).

#### (ii) Coastal upwelling index (monthly, 1997-2016, all sites)

The Bakun index (Bakun, 1973) provides an index of large-scale coastal upwelling and specifically describes the volume of water that is transported offshore from Ekman transport (http://www.pfel.noaa.gov/ - sites = 33N-119W and 39N-125W. This index has been used as a large-scale proxy for processes that may favor larval retention and advection from shore in addition to its value as a predictor of coastal productivity (Menge et al., 2011; Menge & Menge, 2013; Shkedy & Roughgarden, 1997; Wing et al., 2003). We emphasize that this index is not a location specific metric nor does it consider fine-scale nearshore hydrodynamic processes that are also likely to affect larval retention (Fisher et al., 2014; Morgan et al., 2009; Morgan et al., 2016; Morgan et al., 2018; Shanks & Shearman, 2009; Shanks & Eckert, 2005; Shanks & Morgan, 2018; Shanks et al., 2017) which remain outside the scope of this study, but worthy of investigation as drivers of settlement trends. Rather we include the Bakun index as an indicator of broader scale coastal upwelling that may affect regional trends in larval supply (e.g. Roughgarden et al., 1988).

#### (iii) Sea surface chlorophyll (monthly, 1997-2016, all sites)

Satellite imagery of sea surface chlorophyll a provides a spatially and temporally resolved estimate of phytoplankton biomass (mg m^-3^) that is not available from in situ sampling. We used version 3.1 of the OC-CCI merged ocean color time series (Sathyendranath et al., 2018) that combines SeaWIFs, MERIS, MODIS and VIIRS to provide the temporal and spatial coverage required for this study. For each larval settlement collection site, we aggregated data into a 30-day moving average (30 days prior to brush collection) for an area 10 km from shore that stretched 150 km alongshore (the average Lagrangian estimate for dispersal distances for species with a planktonic larval duration (PLD) of 30-days (but see Shanks, 2009; Siegel et al., 2003)). For the site at Anacapa Island, we included any point within a 150 km radius and within 10 km of any coastline. We used 30-day moving averages for chlorophyll and temperature (below) because larvae were exposed to these conditions for at least 30 days prior to settlement, and because averaging over 30 days minimized the effects of serial autocorrelation in the data.

#### (iv) Sea surface temperature (monthly, 1997-2016, all sites)

We used the 30-day moving average of sea surface temperature, derived from Pathfinder AVHRR (Reynolds et al., 2007) (advanced very high resolution radiometer) that was optimally interpolated at daily and 0.25 degree latitude/longitude resolution. We spatially aggregated data in the same way as sea surface chlorophyll.

#### (v) Fall kelp canopy biomass (annual, 1996-2016, Santa Barbara Channel & San Diego only)

Giant kelp (*Macrocystis pyrifera*) is a preferred food and a major constituent of *S. purpuratus* diets in southern California (Foster et al., 2015). The regional biomass of giant kelp *Macrocystis pyrifera* can fluctuate dramatically from year to year (Bell et al., 2015) and cause orders of magnitude variations in *S. purpuratus* fecundity (Okamoto, 2014). We estimated intra and interannual variability in the biomass of giant kelp from Landsat Thematic Mapper satellite imagery (Bell, 2017). Data were aggregated for the Santa Barbara Channel (including islands and mainland from Point Conception to Santa Monica Bay through Ventura County) or the San Diego region (mainland coast from the US-Mexico through Orange County including Santa Catalina and San Clemente Islands). We used the 3-month running mean of kelp canopy biomass during the period leading into the spawning season (Sept-Nov) because marked declines in reproductive capacity require several months of consistently low food supply (Okamoto, 2014).

#### (vi) Adult sea urchin density in the Channel Islands (annual, 1997-2016, Santa Barbara Channel only)

The abundance of adult sea urchins can also fluctuate over time, which can affect larval production and supply. We used the spatial geometric mean of adult biomass density (kg m^-2^) of purple sea urchins (calculated from surveys of density and size structure from the Channel Islands Kelp Forest Monitoring program (Kushner et al. 2013) and a size-biomass model from the Santa Barbara Coastal LTER (Reed, 2018) as an index of the spawning biomass of purple sea urchins in the Santa Barbara Channel. We used the geometric mean (exponentiated log-scale mean) as it accounts for spatial differences in overall mean abundance among sites. We use a biomass index that incorporates size structure and density instead of just density because individual maximum fecundity increases with size. For details on calculation of biomass from the KFM data see the Electronic Supplement.

We allowed regression coefficients to vary among major regions (Fort Bragg, Santa Barbara, and San Diego sites). We used this level of inference because the covariates are aggregated over spatial regions that encompass all sites within a region due to the long planktonic larval duration of sea urchins. The regression analyses included either all global covariates (ENSO, PDO, NPGO) or all local covariates (temperature, chlorophyll, adult densities, giant kelp biomass, and the Bakun upwelling index). We conducted separate analyses for these two groups of covariates because global indices were directly correlated with local covariates, or in the case of temperature were partially derived from them. For the analysis with local covariates we isolated data to 1997 and onward to accommodate the range of chlorophyll data availability, while global analysis included all data. Thus, our analyses provided non-mechanistic explanatory variables for global covariates and mechanistic explanatory variables for local covariates.

## Results

### Spatio-temporal trends in larval settlement

Substantial interannual variability in larval settlement of *S. purpuratus* was observed among years (Figure 3, Appendix Figure S1). Fluctuations in larval settlement were highly synchronous among sites within each of the two regions in the Southern California Bight (r = 0.73 and 0.85 for sites within the Santa Barbara Channel and San Diego, respectively; Figure 3b, c). Within the Santa Barbara Channel, pairwise correlations in interannual trends involving Gaviota, Ellwood and Stearns Wharf were higher (r = 0.86 to 0.90) than those involving Anacapa (r = 0.46 to 0.71). Anacapa began to decline in 2012, while the declines at the other sites did not begin until 2014 (Figure 3b, Appendix Figure S.1 b-e). While all of the Santa Barbara Channel sites except Anacapa exhibited eight continuous years of above average settlement following the low in 2005, San Diego sites remained mostly below average after 2003(Figure 3c, Appendix Figure S.1 f-g). This trend produced modest synchrony among San Diego and Santa Barbara sites from 1991 through 2007 (mean r = 0.68), but none thereafter (mean r = -0.06). In contrast, interannual trends in larval settlement at Fort Bragg were largely uncorrelated throughout the time series with sites in the Santa Barbara Channel (Figure 3a vs. 3b, mean r = -0.23) and sites in San Diego (Figure 3a vs. 3c; mean r = 0.12).

**Figure 3:**
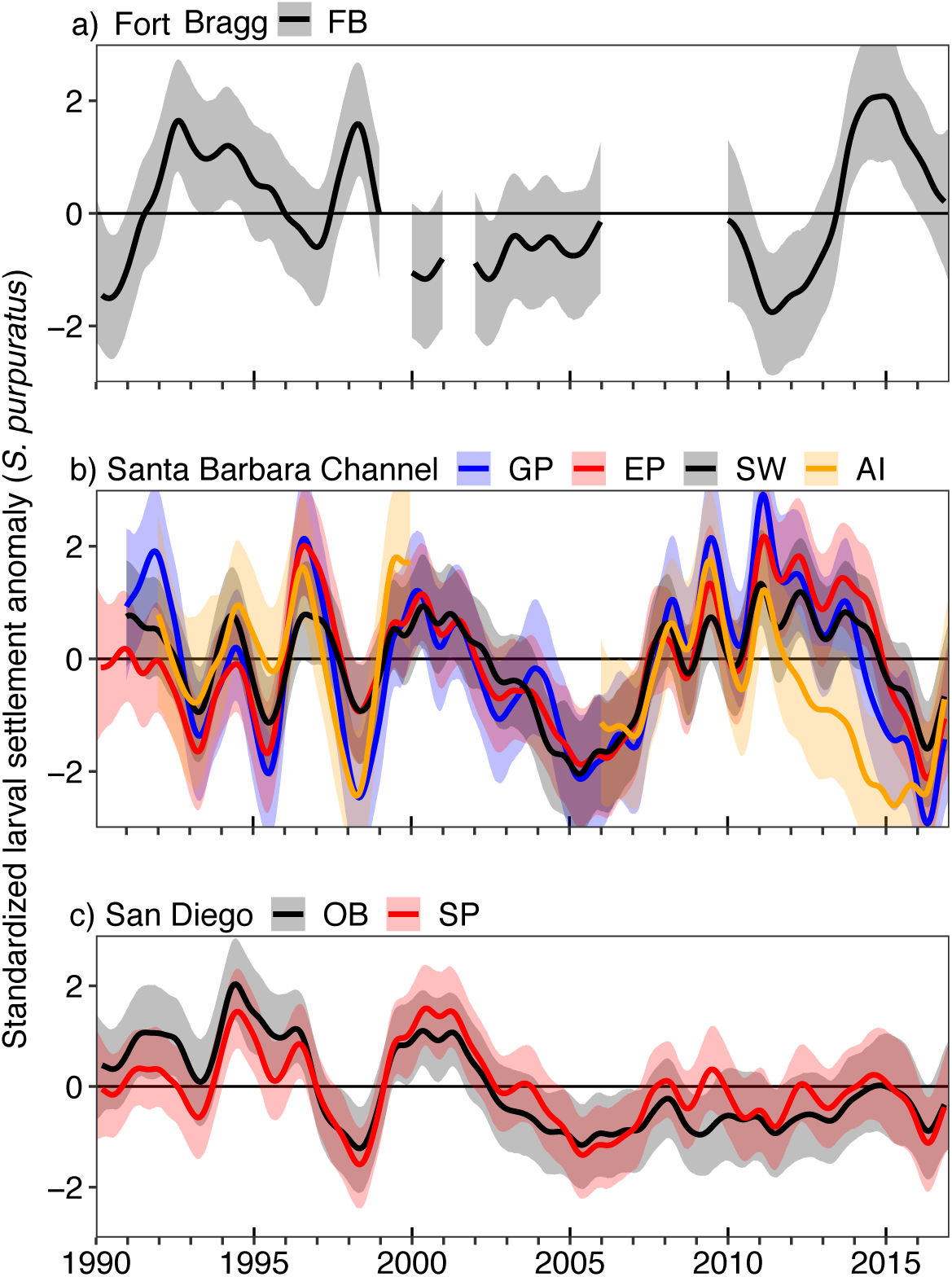
Standardized annual scale trends (log-scale, standardized to one SD) of larval settlement *S. purpuratus* from 1990 through 2016 for: (a) Fort Bragg, (b) sites in the Santa Barbara Channel, and (c) sites in San Diego. Colors within represent individual sites associated with the legend. Interannual trends are estimated using a 3/2 Matérn Gaussian process covariance within a Bayesian multivariate state-space model accounting for spatiotemporal correlations.

Larval settlement in southern California (San Diego and the Santa Barbara Channel) was highly seasonal, with similar patterns among sites (Figure 4a, Appendix Figure S2). On average 90% of recruitment occurred from March to July with a single peak in late April/early May (Figure 4a, Appendix Figure S2), and by July settlement, on average, was an order of magnitude lower than during the April-May peak, and by September is two orders of magnitude lower. By contrast, recruitment at Fort Bragg in northern California extended over a longer period of time (90% occurred, between January and September) and typically included two peaks per year (a large peak around July and a smaller peak in February and March; Figure 4a). The seasonal peaks in settlement in southern California coincided with peaks in sea surface chlorophyll (Figure 4a vs. 4b.) and troughs in sea surface temperature (Figure 4a vs. 4c). At Fort Bragg in northern California the primary peak occurred slightly after the peak in chlorophyll a (Figure 4a. vs. 4b). These peaks in settlement were largely consistent among years, albeit with large variability in magnitude (Shown by red lines in Appendix Figure S1) and some interannual variability within sites (see Appendix Figure S3 for a comparison in seasonal trend among years).

**Figure 4:**
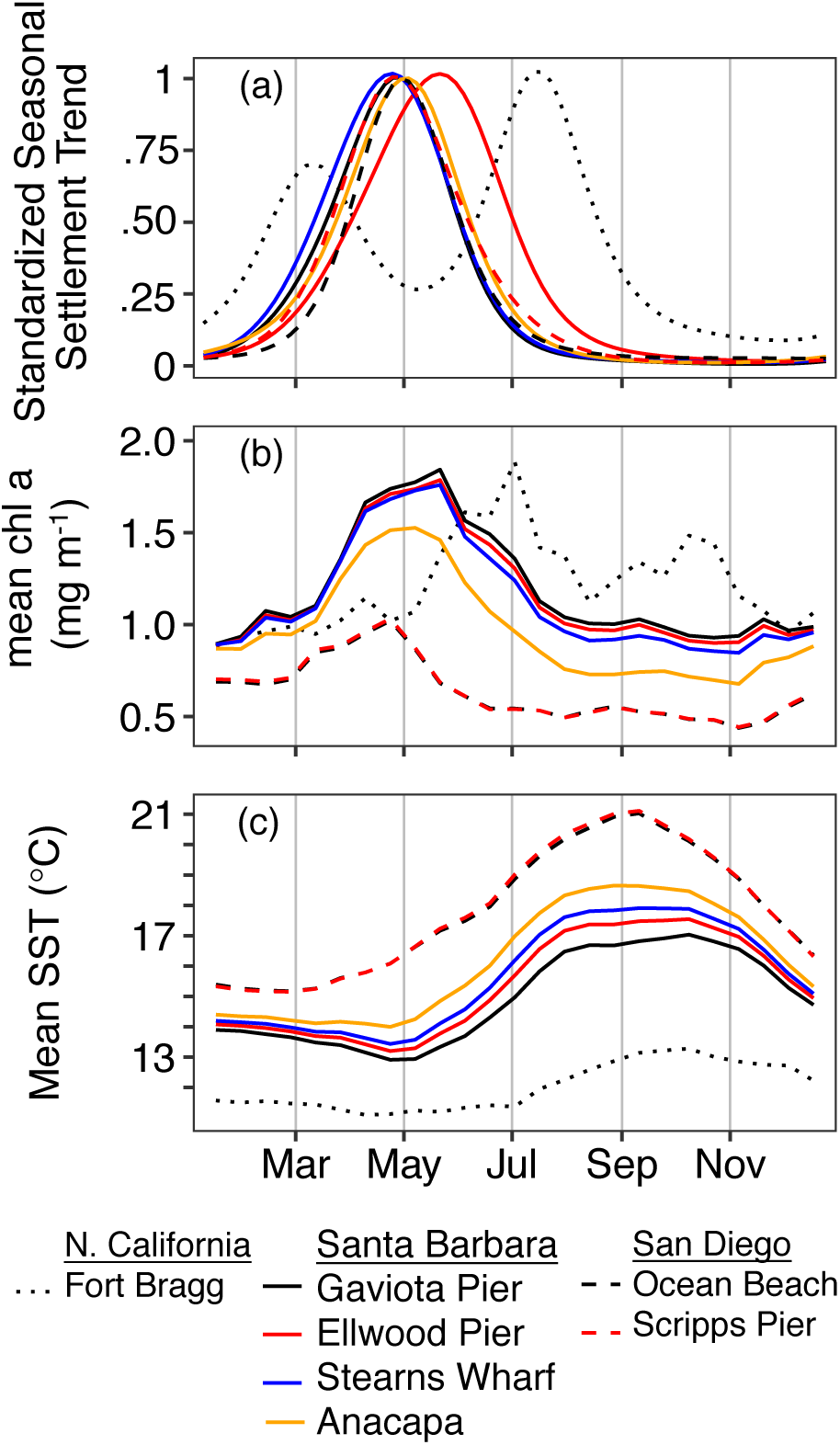
Seasonal trends at each site for: (a) standardized larval settlement of *S. purpuratus*, (b) biweekly mean sea surface chlorophyll a (chl a) and (c) biweekly mean sea surface temperature (SST). The seasonal trend in larval settlement at each site was standardized by the mean maximum seasonal value. Chl a and SST were measured using satellite imagery within 25 km of the coastline and 150 alongshore km of the location where settlement was sampled (hatched areas in Figure 1). See Methods for details.

### Relationship between benthic year-class strength and larval settlement

Recruitment of juvenile purple urchins at shallow subtidal reefs in the Santa Barbara Channel exhibited a significant, positive correlation with larval settlement to brushes two years prior (Figure 5, negative binomial GLMM – 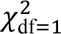 = 9.48, p = 0.002). These results are robust to whether collection sites included only Anacapa (the sole site in the Channel Islands) or incorporated serial correlation using a site-specific first-order autoregressive model in the error covariance matrix. Years with the highest larval settlement corresponded with a nearly three-fold increase in the mean density of juvenile urchins two summers later (Figure 5). Over the time series, average settlement densities in the Santa Barbara Channel varied by more than three orders of magnitude among years. This interannual variation in settlement (averaged across the sites in the Santa Barbara Channel) corresponded to more than a three-fold increase in the density of benthic juveniles on average.

**Figure 5:**
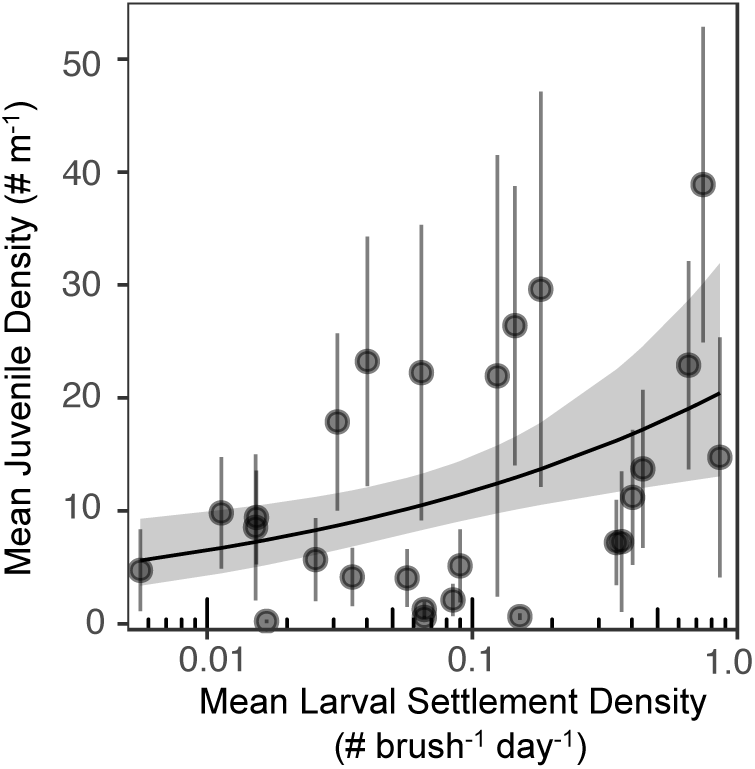
The relationship between the summer density of juvenile purple sea urchins on reefs in the Santa Barbara Channel and the density of recruits on larval collectors during March-July of the previous year (i.e., 12-20 months prior). Points and error bars represent annual means (+/-SE) averaged over all sites. The line with the band represents the estimated relationship and 95% uncertainty interval.

### Relationships between larval settlement and biotic and abiotic conditions

Larval settlement of purple urchins showed strong correlations with local sea surface temperature (SST, Figure 6 a-g, Figure 7 a) as well as the major climate indices (e.g. ENSO - Fig 6 h-n, Figure 7f). For temperature and ENSO, the sign of the correlation at Fort Bragg was opposite that at sites in the Santa Barbara Channel and San Diego. The correlation between settlement and temperature varied from strongly negative at sites in San Diego and Santa Barbara to positive at the Fort Bragg site. Correlations between larval settlement and adult urchin density, upwelling, chlorophyll and kelp biomass were either not different from zero or negative, which was opposite of that expected at all sites (Figure 7 b-e). Larval settlement and sea surface chlorophyll were uncorrelated at Fort Bragg and in the Santa Barbara Channel and negatively correlated in San Diego – the opposite of the hypothesized relationship that more food in the plankton would lead to higher larval settlement (Figure 7 d). The network of correlations among these variables are depicted in (Appendix Figure S.3).

**Figure 6:**
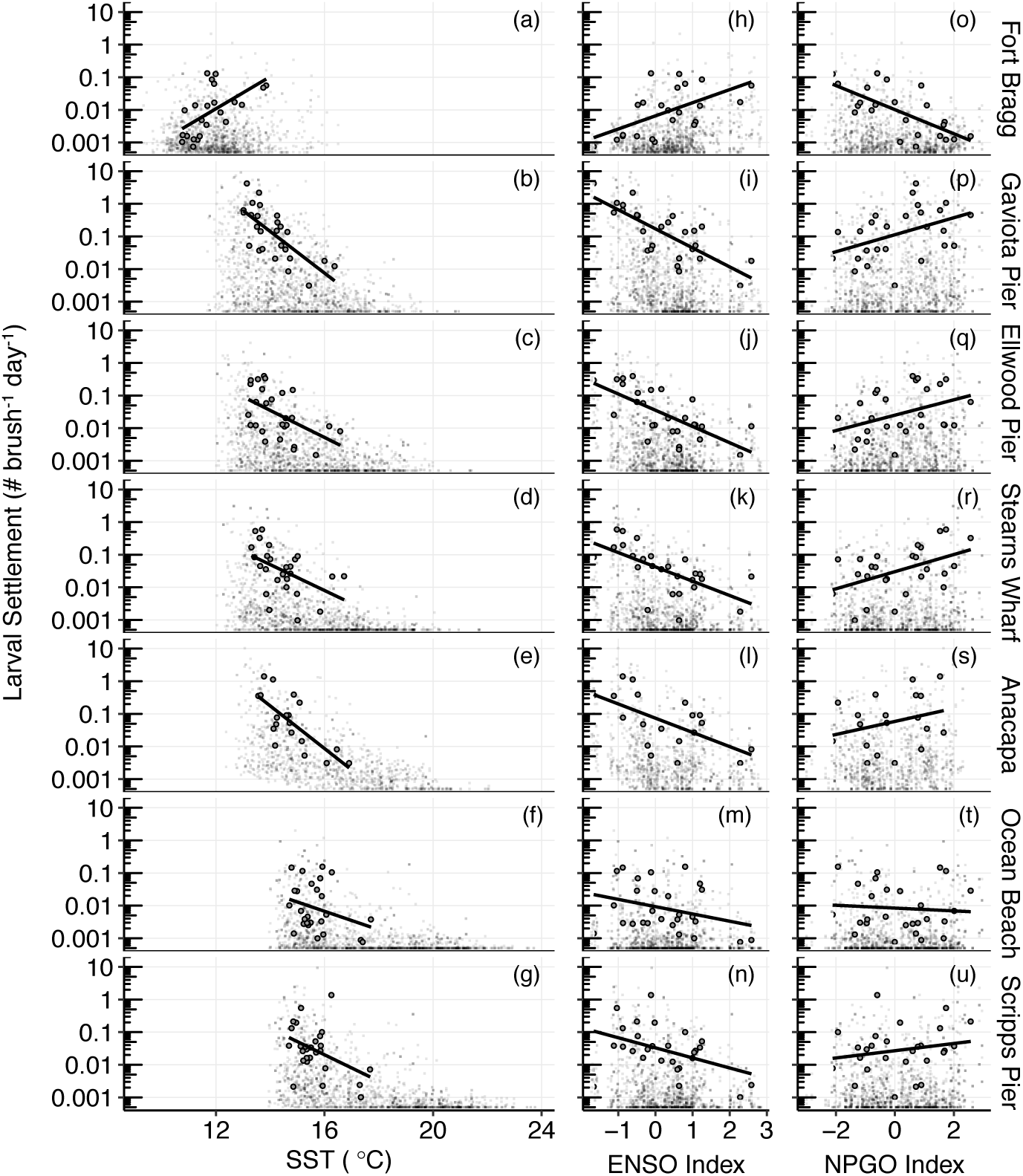
Standardized coefficients for the relationships between larval settlement of purple sea urchins at each site and: (a-g) sea surface temperature, (h-n) the multivariate ENSO index, and (m-u) the North Pacific Gyre Oscillation (NPGO). Small grey points represent biweekly means and black points and lines represent annual means.

**Figure 7:**
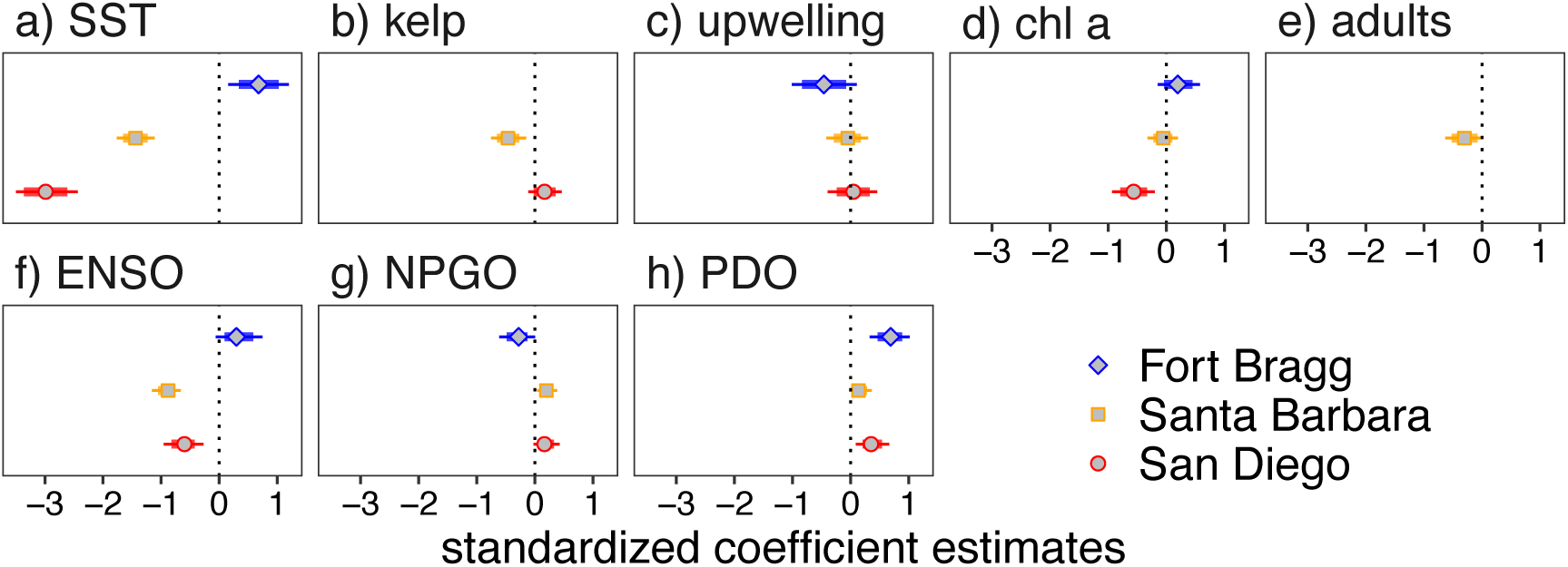
Standardized Bayesian multiple regression coefficients for relationships between *S. purpuratus* settlement and local covariates (a-e) or global climatic indices (f-h). Models with global and local variables were fit separately. Coefficients were estimated using a multiple regression nested within a Bayesian multivariate time series model. Coefficients are *a priori* biased towards zero using a regularized horseshoe prior for variable selection. For details see Methods, Figure 2 and appendix for details.

Larval settlement in southern California was orders of magnitude lower during warm, El Niño conditions and during the negative phase of the NPGO with the more southern sites in San Diego responding more strongly to temperature than those further north in the Santa Barbara Channel (Figure 7 a). In contrast, Fort Bragg, the northern most site, responded positively to warmer El Niño conditions, negatively to the NPGO, and positively to the PDO. Importantly, relationships observed between larval settlement and ENSO and temperature in southern California were opposite of those observed at Fort Bragg (Figure 7 f, g). The correlations between larval settlement and SST and ENSO occurred on roughly 3-5 year cycles, while the correlation with NGPO and PDO occurred on decadal scales.

## Discussion

Climate variability can significantly impact marine ecosystems by affecting recruitment, which in turn influences the dynamics of populations and communities. Yet determining which species and locations will respond to shifting climate remains a difficult task. Using a multi-decadal, high frequency, and spatially extensive time series we show how large-scale climatic variation can give rise to spatially different responses in the settlement of larval sea urchins. We found that settlement patterns were synchronous within Santa Barbara and San Diego and synchronous between these two regions until the late 2000s. Importantly, these sites responded negatively to elevated temperature and ENSO events. This contrasted sharply with settlement at our northern California site (Fort Bragg), which was positively correlated to temperature and ENSO. This difference was most obvious during the two strongest ENSO events in 1998 and 2014, when settlement responded dramatically in opposite directions.

While many recruitment-environment correlations often break down over time (Myers, 1998), the correlations between oceanographic factors and larval settlement in our study were evident over a 27-year period that included multiple major ENSO events and several minor ones. We observed repeatable patterns of diminished larval settlement in response to temperature and ENSO events in southern California with opposite responses at Fort Bragg in northern California. Others have found larval settlement in sea urchins to be positively or negatively related to temperature depending on the species (Ebert, 1983; Hernández et al., 2010; Himmelman, 1986). Our study is among the first to show that the relationship between larval settlement and temperature can vary dramatically within a species depending on location. Although sea urchin larval settlement data at the fine temporal scale of our study are not available prior to 1990, there is a history of evidence for ENSO-related recruitment failures in sea urchin populations of southern California. Between 1969 and the early 1980’s, recruitment of juvenile sea urchins was anomalously low during El Niño years (Ebert, 1983; Tegner & Dayton, 1991). How patterns of larval settlement change in the future will undoubtedly depend on the processes underlying their associations with ENSO events.

Climate related fluctuations in the supply of larvae have been hypothesized to affect coastal species around the world (Caley et al., 1996; Grosberg & Levitan, 1992; Underwood & Fairweather, 1989). In the case of sea urchins, larval supply has been implicated for dramatic changes in the state of benthic communities (Estes & Duggins, 1995; Hernández et al., 2010; Ling et al., 2009). Yet empirically demonstrating such links has historically proved challenging because recruitment can be attenuated by high mortality of newly settled larvae (Connell, 1985; Rowley, 1989), increasing the need for high frequency data of larval settlement spanning multiple years. Data such as ours that meet these criteria are rare.

The potential factors affecting larval supply that we examined other than temperature and climate showed no meaningful correlations with larval settlement. Of particular note was the lack of a correlation with ocean chlorophyll revealed by our analyses. This finding differed from that observed for the tropical sea urchin *Diadema aff. antillarum* (Hernández *et al*. 2010) and other echinoderms (e.g. Fabricius et al., 2010; Lamare & Barker, 1999) and highlights a need for exploring mechanistically how spatial and temporal variability in ocean circulation and phytoplankton productivity affect patterns of larval settlement. These outcomes do not suggest that food is not important for urchin larvae, only that interannual variation therein did not explain the interannual trends in settlement. Indeed, the mean seasonal trends in both southern and northern California (which we show are synchronous in southern California) corresponded to peaks in primary productivity, and both are regions of high productivity (Kahru et al., 2012).

Similar to other studies (Shanks & Shearman, 2009; Shanks & Morgan, 2018) we found no relationship between large-scale fluctuations in indices of upwelling and larval settlement, which runs counter to hypotheses that large-scale coastal upwelling shapes patterns of larval supply by broadly altering productivity and retention (Menge & Menge, 2013; Roughgarden et al., 1991). The major ocean currents in our study system (the Southern California Counter Current and the California Current) are characterized by stochastic and spatially variable eddies, fronts, filaments and bores (e.g. Bassin et al., 2005; Davis et al., 2008; DiGiacomo & Holt, 2001), whereas flows closer to shore tend to promote larval retention along wave exposed shores (Morgan et al., 2016; Shanks et al., 2017). As a result, offshore transport and retention off the coast of California is spatially nuanced and temporally inconsistent (Shanks & Eckert, 2005). Such heterogeneity not only influences larval transport, but also when and where larvae encounter productive food environments. Consequently, if larval food limitation plays a role in shaping the dynamics of urchin settlement at our sites, then it likely arises out of more complex dynamics than our spatially aggregated composite metrics were able to detect. For example, processes that shape stratification, front formation and spatially isolated phytoplankton blooms in combination with larval behavior may play a more important role than predicted by simplifications of the higher or lower spatially averaged phytoplankton biomass or greater or lesser coastal upwelling.

The oceanographic processes that affect spatial distribution and survival of larvae may also affect predation driven mortality by altering the densities of predators and their larval prey. While predation on echinoplutei certainly occurs, its relative importance in shaping larval supply to the benthos remains difficult to quantify (Rumrill, 1990; Vaughn & Allen, 2010). For *S. purpuratus* plutei, predation may be lower than for other taxa because they contain chemical and structural defenses and display predator avoidance behavior (Cowden et al., 1984), particularly at later stages of development, which are far less susceptible to predation (Rumrill, 1985). This is supported by field studies that have shown predator mortality to be comparatively low for echinoids (Allen & McAlister, 2007; Johnson & Shanks, 2003), which in part may be due to the fact that accounting for natural densities of echinoplutei can dramatically reduce estimates of laboratory predation rates (Johnson & Shanks, 1997). Integrating mechanistic models of ocean circulation and mortality was beyond the scope of our study, yet such approaches are needed to develop a more thorough understanding of processes that control settlement dynamics across large gradients in oceanographic settings as they can control both population dynamics and range limits (Gaylord & Gaines, 2000).

Although climate related changes can alter larval supply via their effects on the production of larvae by adults, we found no relationship between larval settlement and regional adult abundance or adult food (kelp). While fisheries research relies heavily on stock-recruit dynamics, there is continual debate about whether adult dynamics actually control recruitment patterns (Gilbert, 1997; Szuwalski et al., 2015). The lack of a positive correlation between adults and larval settlement that we found for purple sea urchins adds to this debate by showing that high abundances of adult urchins and kelp averaged across large spatial scales did not translate into high larval settlement. We note that spatial structure in the dynamics of kelp and adult urchins may have affected this outcome. For example, modest but uniform densities of adult sea urchins in food rich kelp forests can provide very different outcomes for the production of embryos compared to mosaics of dense adults in barren patches interspersed with forested patches having few adults (Okamoto, 2014). Indeed, previous work has demonstrated that assessing population productivity in spatially structured populations requires a spatially structured analysis to account for processes of population regulation and competition (Chesson, 1996, 1998; Chesson et al., 2005; Thorson et al., 2015). That said, we found no evidence that synchronous fluctuations in settlement arose from broad-scale collapses and proliferations in populations of adult sea urchins or kelp at the appropriate lags and time-scales.

Temperature-related effects on larval production and survival and ENSO-related changes in currents or oceanographic features that transport larvae represent two possible explanations for the spatially opposing correlations between larval settlement and temperature and ENSO that we observed. Many species have upper temperature thresholds beyond which reproduction is impaired that are well below the lethal limits for adults. Temperatures in southern California kelp forests routinely exceed 17°C during ENSO events (Reed et al. 2016), which may adversely affect larval supply of purple sea urchins by limiting gamete production (Basch & Tegner, 2007; Cochran & Engelmann, 1975), fertilization (Schroeder & Battaglia, 1985) larval development and gene expression (Padilla-Gamiño et al., 2013; Runcie et al., 2012; Wong et al., 2018), and larval survival (Azad et al., 2012). Such effects may in part explain why larval settlement at Fort Bragg, where sea surface temperatures rarely exceed 16°C was not negatively correlated with temperature. Unfortunately, there are no long-term datasets on reproductive activity or planktonic larval densities and distributions to evaluate if and how the interannual and seasonal trends observed here correspond to variation in reproductive activity and transport of larvae.

Regional circulation is considered to be an important driver of larval supply for many species, including those in the Southern California Bight (Blanchette et al., 2006; Broitman et al., 2005; Hereu et al., 2004; McManus & Woodson, 2012; Woodson et al., 2012). ENSO events can cause major changes in patterns of ocean circulation in the Southern California Bight and elsewhere along the US west coast (Lynn & Bograd, 2002; Mitarai et al., 2009; Pineda & Reyns, 2017). Spatial variability in such patterns could explain the opposing climatic responses in larval settlement between Fort Bragg and our sites in southern California. El Niño events in southern California are associated with lower stratification and reduced internal waves that promote the onshore transport of larvae (Pineda, 1994; Pineda et al., 2018; Shanks, 1983). In contrast, El Nino events in northern California are often associated with relaxed upwelling, downwelling Kelvin waves and increased stratification (Chavez et al., 2002). Although we cannot definitively partition out the effects of climate associated changes in temperature and circulation on recruitment, our observations of larval settlement and subsequent year-class strength point strongly to broader-scale ocean climate effects.

Our finding that larval settlement was a good predictor of regional year class strength in natural populations indicates that large fluctuations in larval settlement of this prominent herbivore could have far reaching ecological impacts that resonate throughout marine ecosystems across a broad geographic region (Pearse, 2006). For example, the high settlement that we observed at Fort Bragg in 2013-2015 coupled with a period of anomalously warm water may explain the marked increase in the abundance of purple urchins and coincident loss of canopy forming kelps reported for northern California beginning in 2014 (Rogers-Bennett & Catton, 2019). Although regional processes appear to drive broad-scale patterns of larval settlement in sea urchins, local hydrodynamics and other factors contribute to considerable fine-scale spatial variation in larval supply and settlement in sea urchins (Ebert et al., 1994; Farina et al., 2018; Hereu et al., 2004; Schroeter et al., 1996; Schroeter et al., 2009). For example, pelagic larval abundance and settlement of *Paracentrotus lividus* varies substantially in space (Hereu et al., 2004; Prado et al., 2012) as does post-settlement survival and abundance due to localized and habitat dependent effects (Boada et al., 2018; Prado et al., 2012). Our inferences focused on temporal trends and comparison of regional-scale patterns, rather than trying to infer causes and consequences of variation in average settlement in space. Despite these caveats, our analyses show remarkable synchrony in settlement within southern California and strong association with regional benthic recruitment in the Santa Barbara Channel. Collectively, our results suggest that predicted changes in ocean climate that lead to more frequent or severe marine heatwaves (Frölicher and Laufkötter 2018) will reduce purple urchin recruitment in southern California, but increase it in northern California with potentially cascading effects on benthic ecosystems similar to those recently observed in northern California.

Oceans are experiencing simultaneous shifts in temperature, water chemistry, productivity and circulation, which have profound consequences for the ecological structure and function of marine systems and the services that they provide (Wong et al., 2014). Our findings provide valuable insights into the ecological consequences of climate related effects on patterns of larval settlement of an important reef herbivore whose distribution spans most of the Pacific coast of North America. Future investigations aimed at determining the specific biotic and abiotic processes that regulate larval settlement in ecological important species such as *S. purpuratus* will undoubtedly improve our understanding of how climate fluctuations affect regional population dynamics to alter the structure and functions of marine ecosystems.

## Acknowledgements

Funding was provided by the California Urchin Commission, the South Bay Cable Committee, the California Department of Fish and Wildlife Director’s Sea Urchin Advisory Committee, and the National Science Foundation’s support of the Santa Barbara Coastal Long Term Ecological Research program. We are indebted to Tom Ebert and John Dixon, who initiated the larval settlement time series with SS. The Channel Islands Naturalist Corps volunteers, Channel Islands National Park Staff, and Channel Islands long term Kelp Forest Monitoring Program were and remain instrumental in collecting settlement data from Anacapa and benthic data in the Channel Islands. John Richards, Clint Nelson, Shannon Harrer, Jenny Wolf, Peter Kalvass, Brigitte Bondoux and many students assisted with field collections, laboratory sampling and data compilation. Sally Holbrook, Russ Schmitt and Cherie Briggs contributed discussions during initial development of analyses. Kyle Cavanaugh, Dave Siegel and Tom Bell provided valuable discussions about giant kelp canopy data. Kyle Cavanaugh and Rachel Simons gave important comments on previous manuscript drafts.

## Supplementary Information - Appendices for

### Estimation of juvenile urchin densities

The KFM program surveys purple sea urchins at the Santa Barbara Channel Islands by counting the total number of individuals in defined areas and measuring approximately 100 individuals at each site. Thus, to model density we used the number of juveniles measured divided by the total number of individuals measured, multiplied by the total number of individuals counted, divided by the area surveyed:

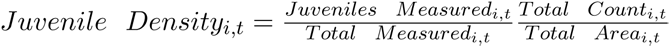

We modeled density in the GLM using the number of measured juvenile urchins (2.5 cm in test diameter-the approximate cutoff size for reproduction - Kenner & Lares 1991). Let represent expected juvenile density at site *i* in year *t*. We constructed the regression as:

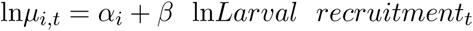

We allowed the intercept (*α_i_*) to vary by site nested within each island because of overall differences in mean juvenile density among sites and islands. We used a negative binomial (NB) with the using direct mean and variance parameterization form:

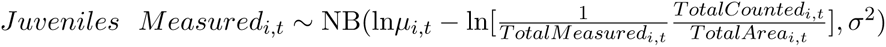

We constructed the model in this format (i.e. with the density denominator in the left hand side of the equation) to maintain the sample size and integer nature of the data while modeling the mean density.

**Figure S.1:**
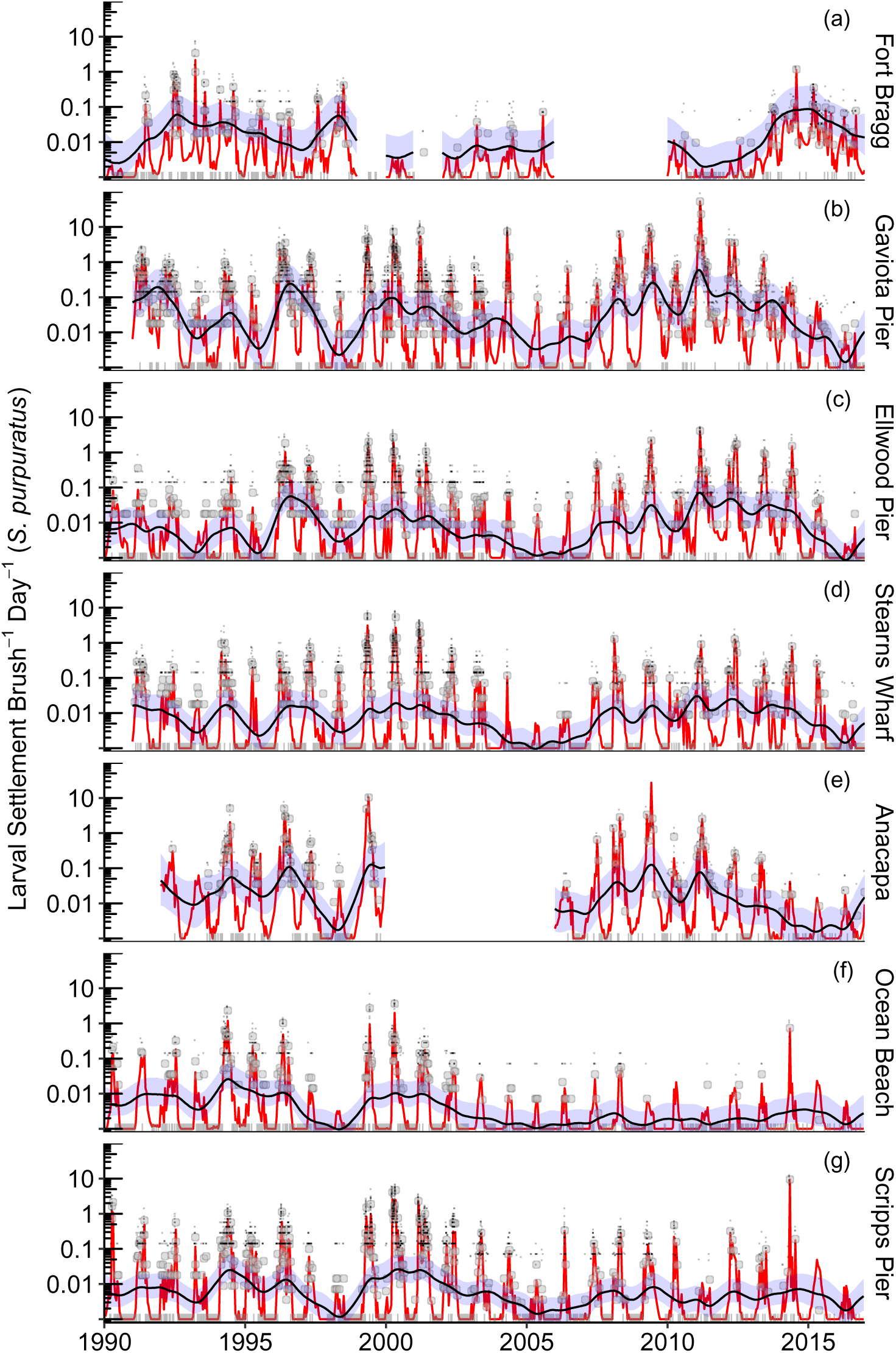
Observed and estimated purple urchin (*S. purpuratus*) settlement trends from 1990-2016 for each site. Large points represent empirical biweekly means, lines on the axis represent biweekly mean values equal to zero, small points represent counts on individual brushes (only positive values shown for clarity, although the models include zero values), red lines represent biweekly estimates, lines with blue 95% uncertainty intervals represent model estimates of the interannual trends. The model estimates the seasonal trend and the interannual trend at each site simultaneously. For reference, interannual trends for *S. purpuratus* shown here are the unstandardized versions of those shown in Figure 2.

**Figure S.2:**
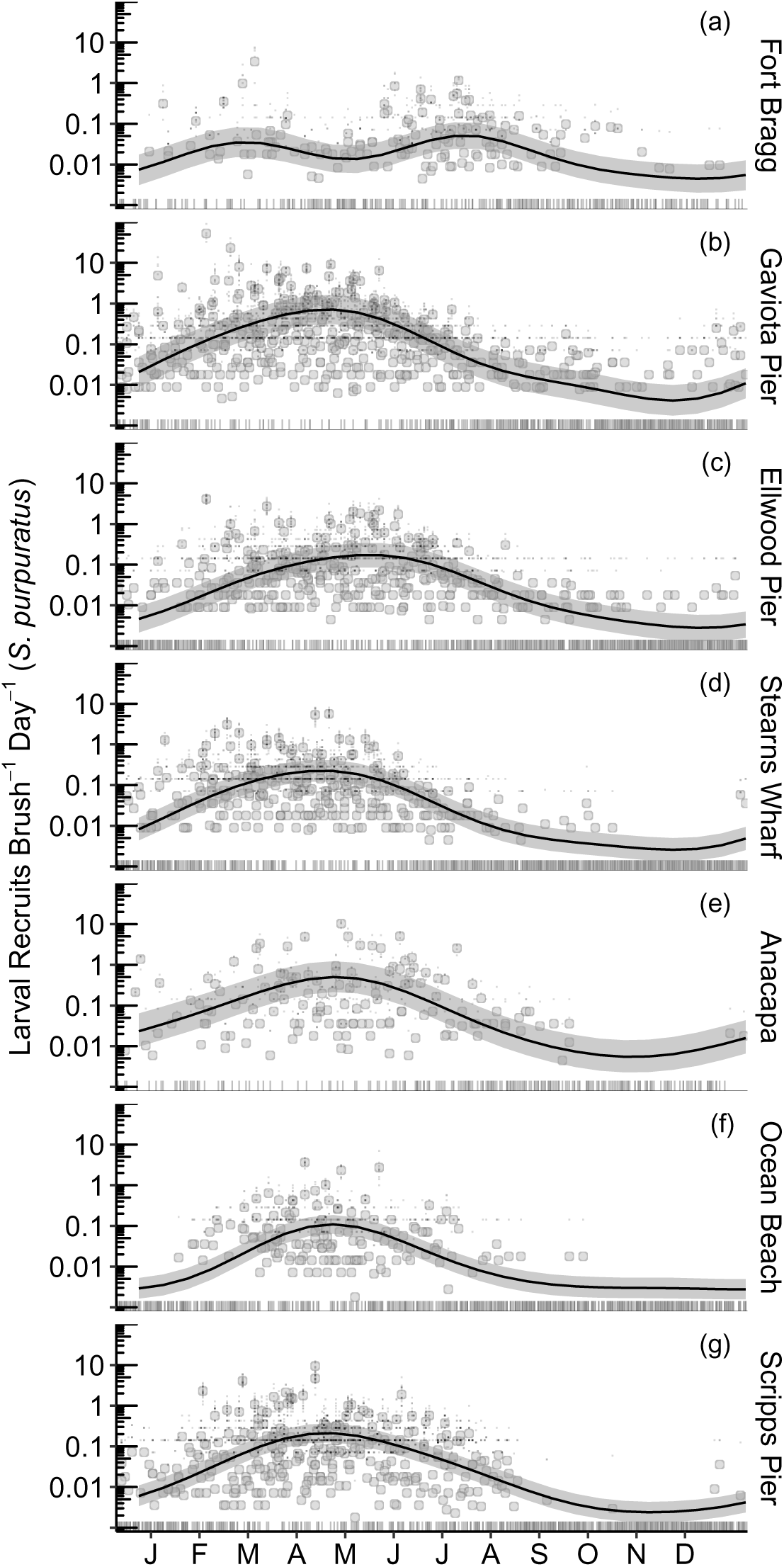
Seasonal settlement trends from 1990-2016 with the mean estimated seasonal trend for each site. Large points represent empirical biweekly means lines on the axis represent biweekly mean values equal to zero, small points represent counts on individual brushes (only positive values shown for clarity, although the models include zero values), black lines represent the posterior mean and bands represent the 95% uncertainty interval. The model estimates the seasonal trend and the interannual trend at each site simultaneously. For reference, seasonal trends for *S. purpuratus* shown here are the unstandardized versions of those shown in Figure 3.

**Figure S.3:**
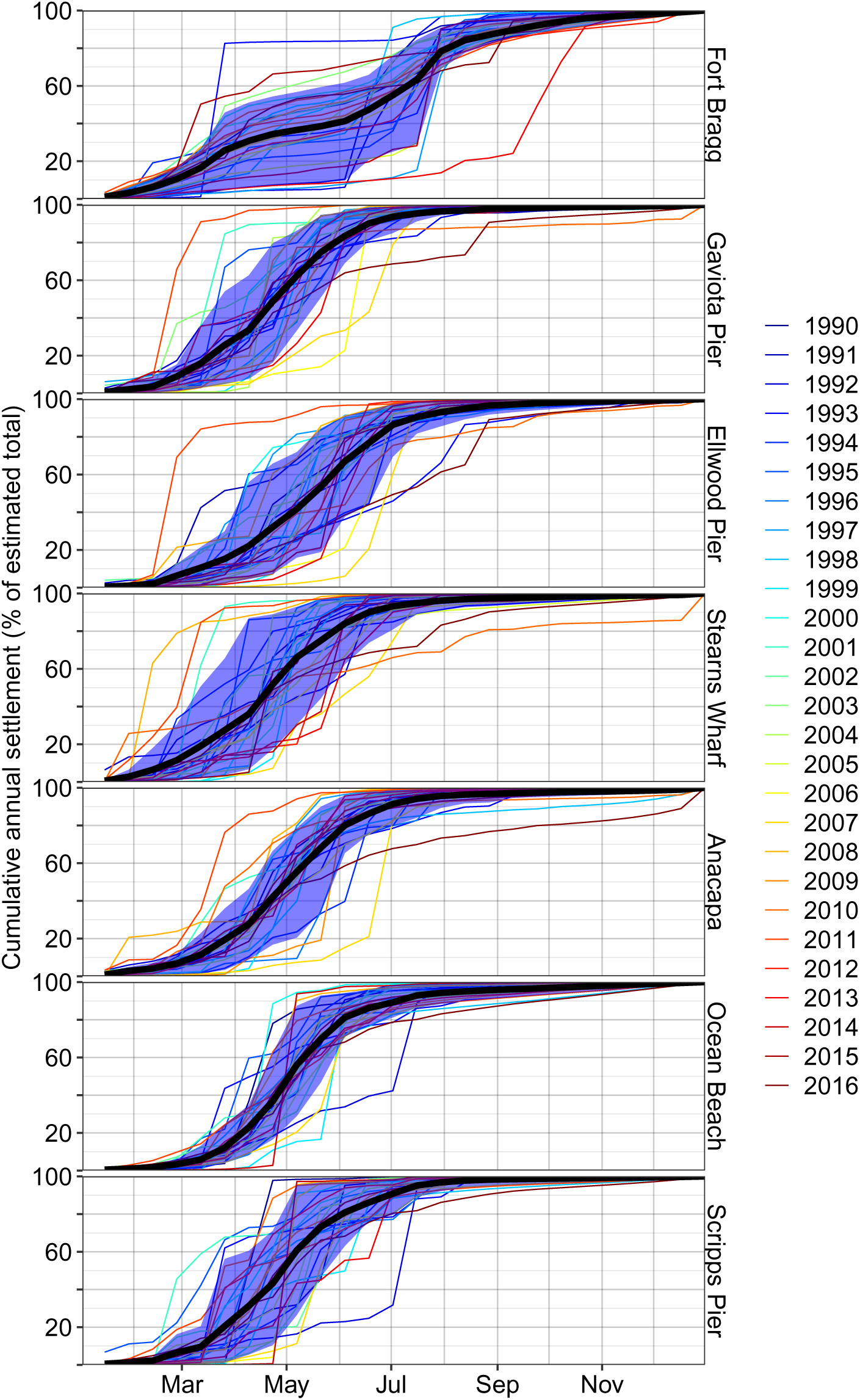
Cumulative annual mean settlement from the estimated biweekly means for each year. Thick black lines represent the overall mean (i.e. the empirical cumulative mean from Figure 3) and thin, colored lines represent the cumulative annual values of the red lines in Figure S1. Blue bands represent the 10 and 90% quantiles for each biweekly period.

### Variable Correlations

To illustrate the intercorrelated nature of the covariates and responses, we estimated a network model for Gaviota Pier that includes the independent model estimates of *S. purpuratus* settlement and its seasonality. To do so, we first constrained network structure by excluding all nonsensical interactions (i.e. chlorophyll does not cause ENSO events and is thus eliminated a priori) and including all known directional interactions (i.e. estimated seasonality affecting settlement for is forced into the network). Learning of the network skeleton is achieved via the Hill Climbing algorithm using the bnlearn (Scutari 2010) package in R. Note that strong collinearity can result in the weaker correlation being ignored (e.g. ENSO over SST with Larval Recruitment).

**Figure S.4:**
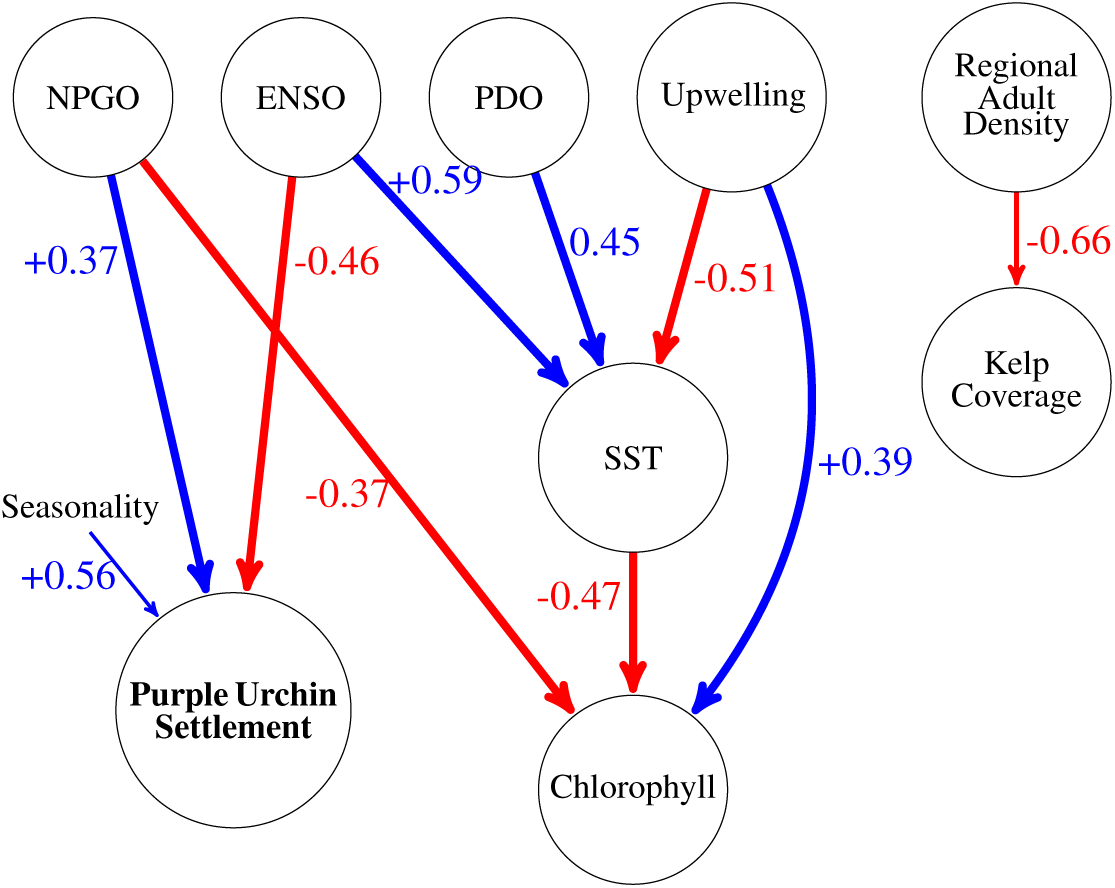
Bayesian network model structure inferred for the Santa Barbara Channel. Numbers represent the partial correlation coefficients accounting for all other variables pointing to that node. All correlations shown are significant at alpha= 0.05

---

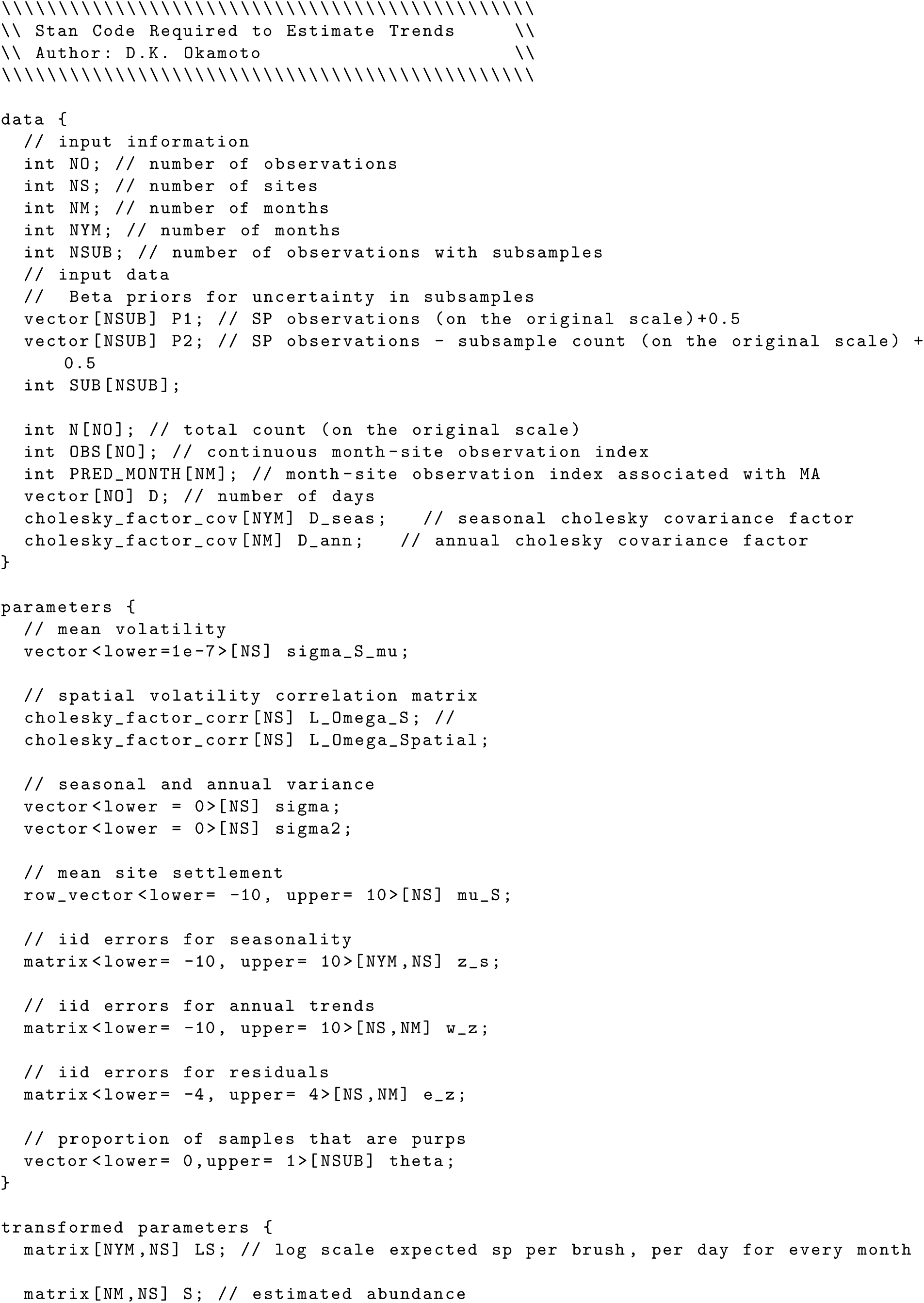
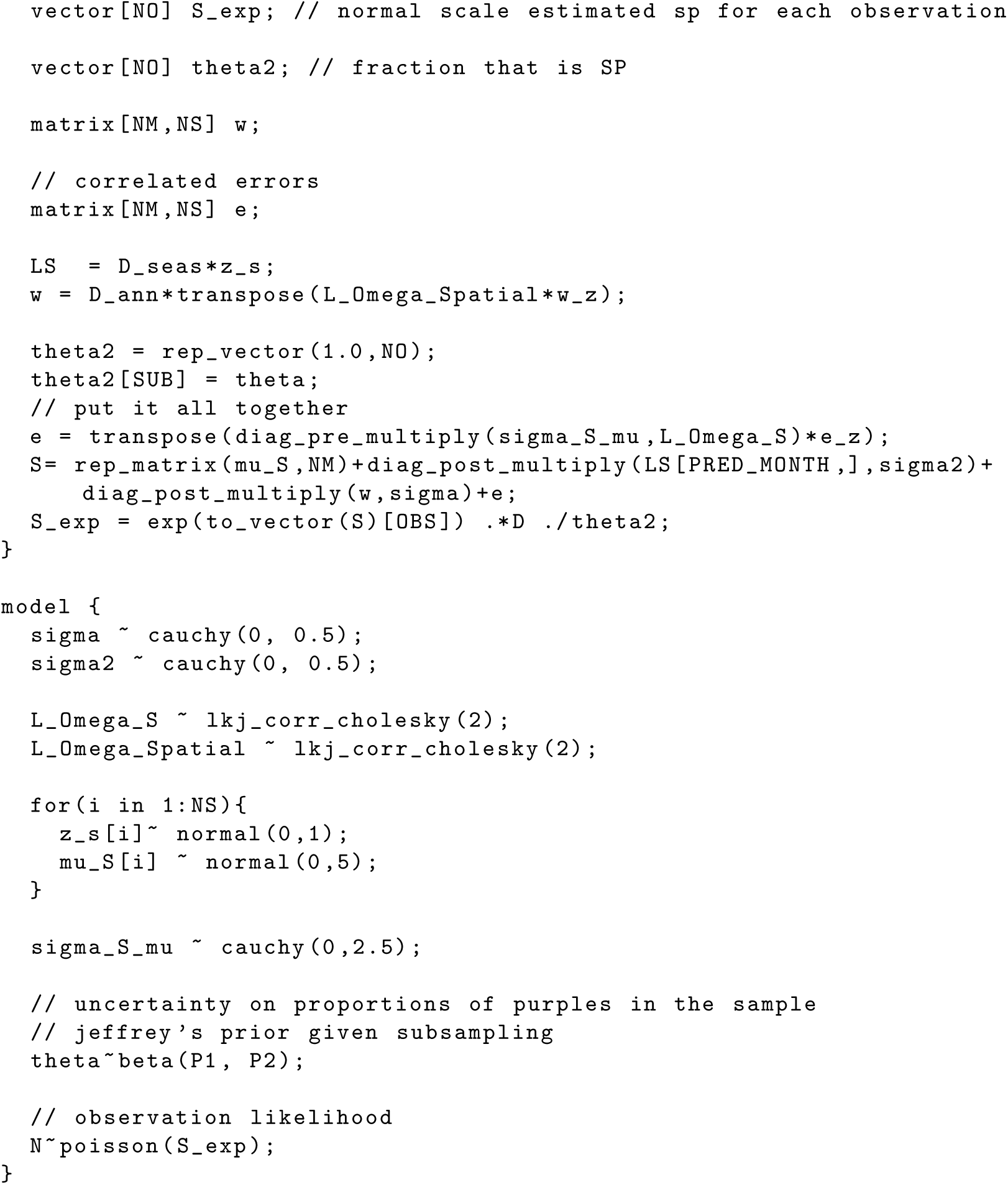

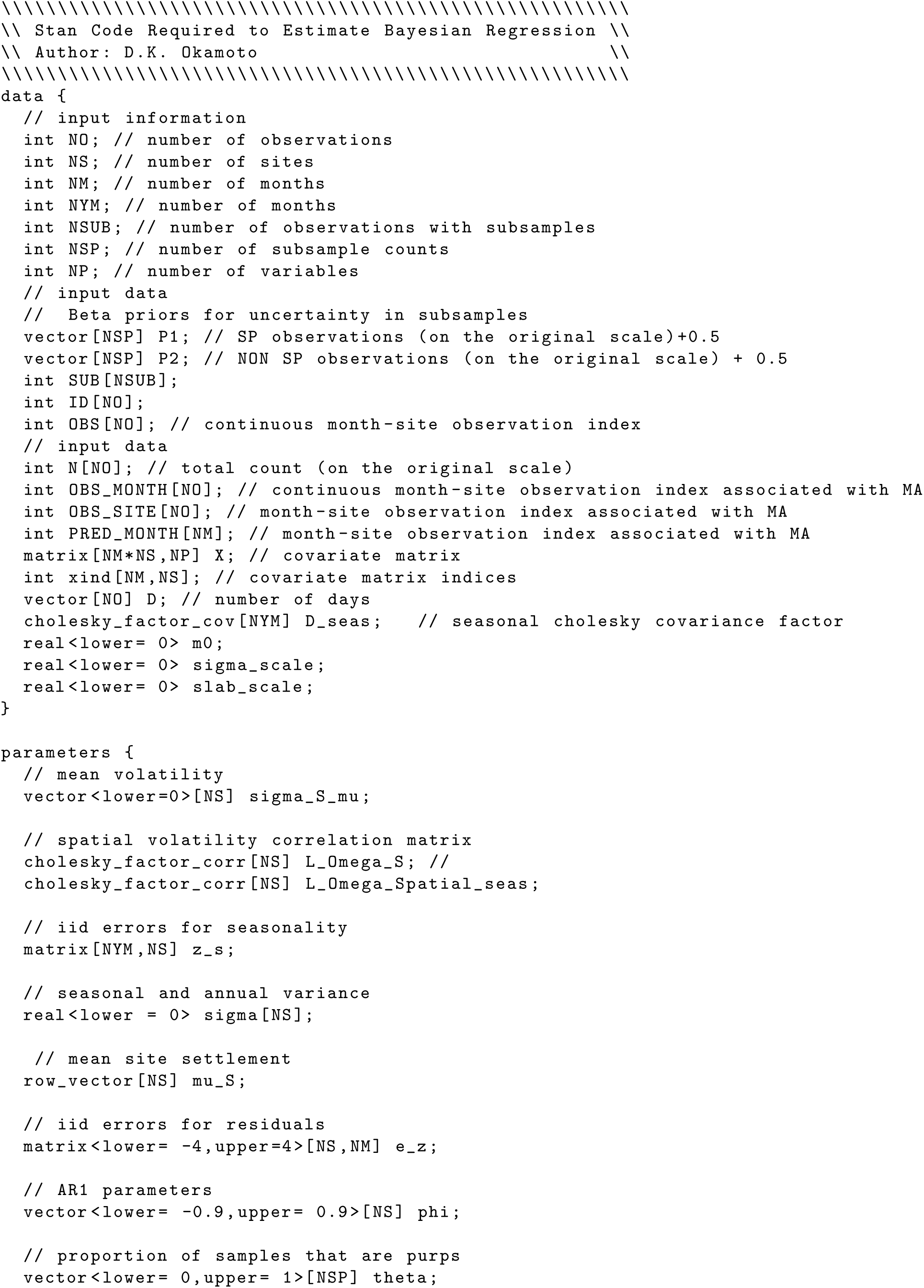
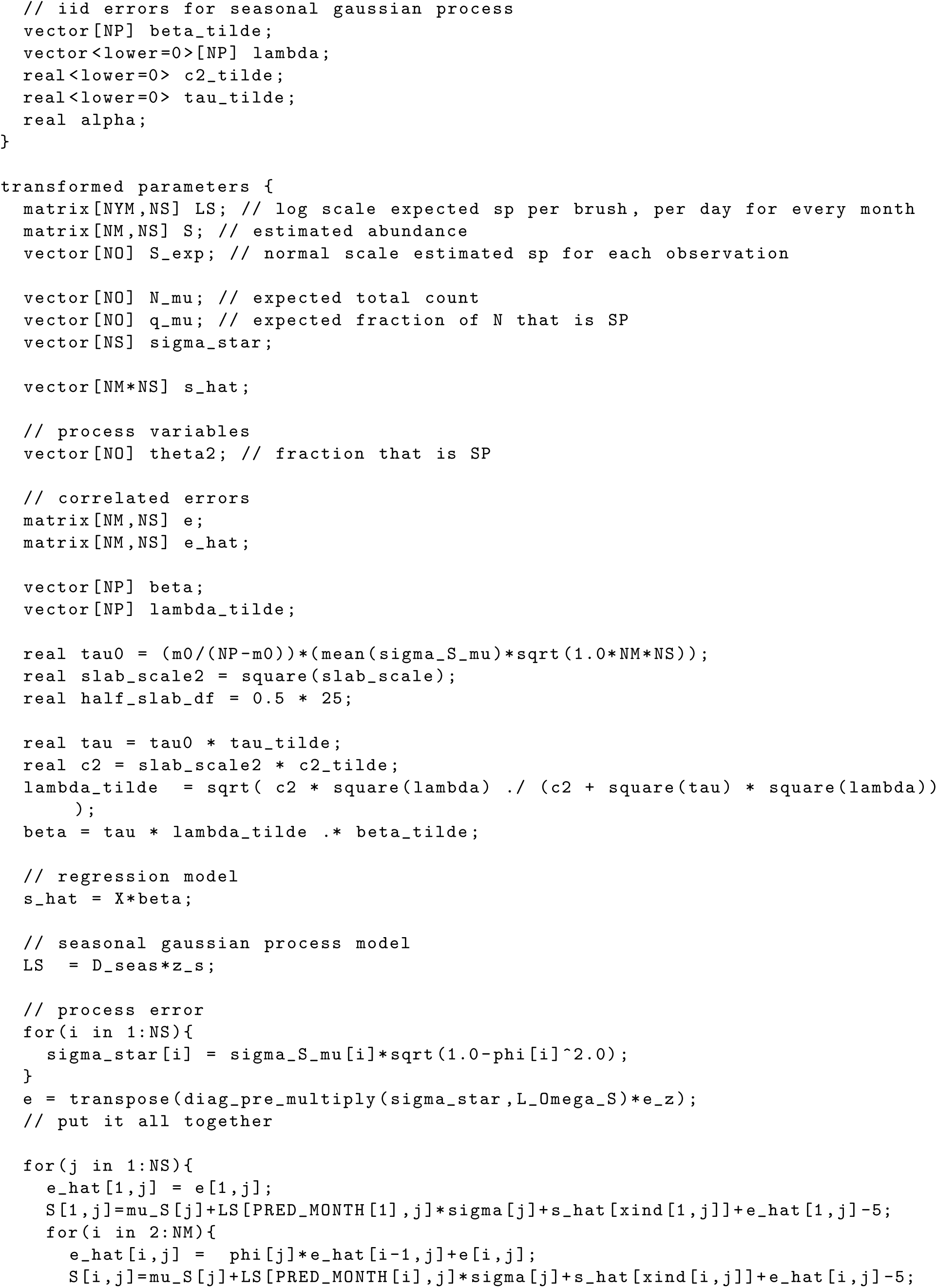
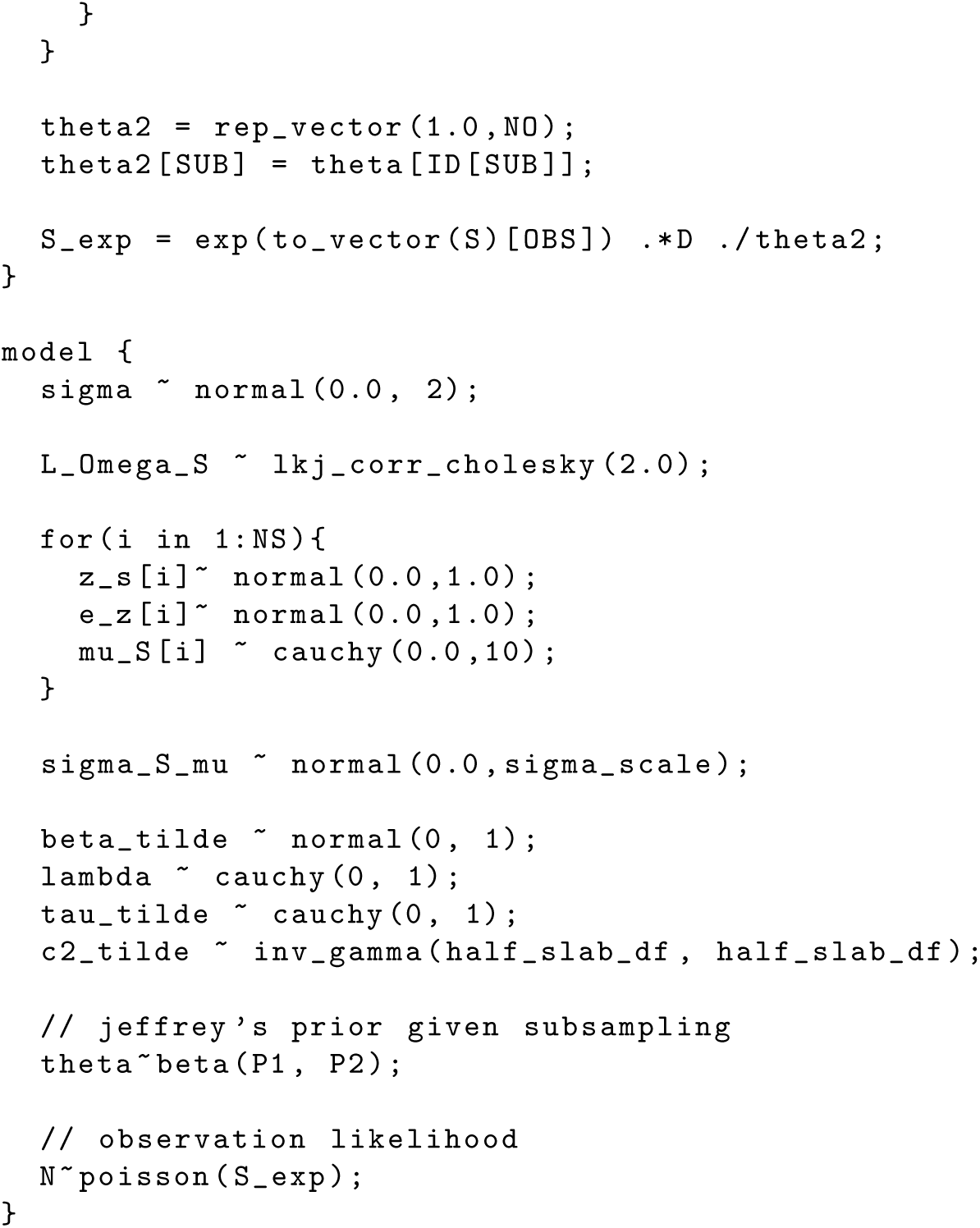

